# Advanced Tissue Technologies of Blood-Brain Barrier Organoids as High Throughput Toxicity Readouts in Drug Development

**DOI:** 10.1101/2024.09.09.611987

**Authors:** Luisa Bell, Claire Simonneau, Chiara Zanini, Elena Kassianidou, Christelle Zundel, Rachel Neff, Bernd Steinhuber, Marco Tecilla, Alex Odermatt, Roberto Villaseñor, Nadine Stokar-Regenscheit

**Author notes:** **Corresponding author:** Nadine Stokar-Regenscheit, Roche Pharma Research and Early Development (pRED), Pharmaceutical Sciences, Roche Innovation Center Basel, Grenzacherstrasse 124, 4070 Basel, Switzerland.

## Abstract

Recent advancements in engineering Complex *in vitro* models (CIVMs) such as Blood-brain barrier (BBB) organoids offer promising platforms for preclinical drug testing. However, their application in drug development, and especially for the regulatory purposes of toxicity assessment, requires robust and reproducible techniques. Here, we developed an adapted set of orthogonal image-based tissue methods including hematoxylin and eosin staining (HE), immunohistochemistry (IHC), multiplex immunofluorescence (mIF), and Matrix Assisted Laser Desorption/Ionization Mass Spectrometry Imaging (MALDI-MSI) to validate CIVMs for drug toxicity assessments. We developed an artificial intelligence (AI) algorithm to increase the throughput and the reliability of histomorphologic evaluations of apoptosis for *in vitro* toxicity studies. Our data highlight the potential to integrate advanced morphology-based readouts such as histological techniques and digital pathology algorithms for use on CIVMs, as part of a standard preclinical drug development assessment.

**Graphical abstract:** 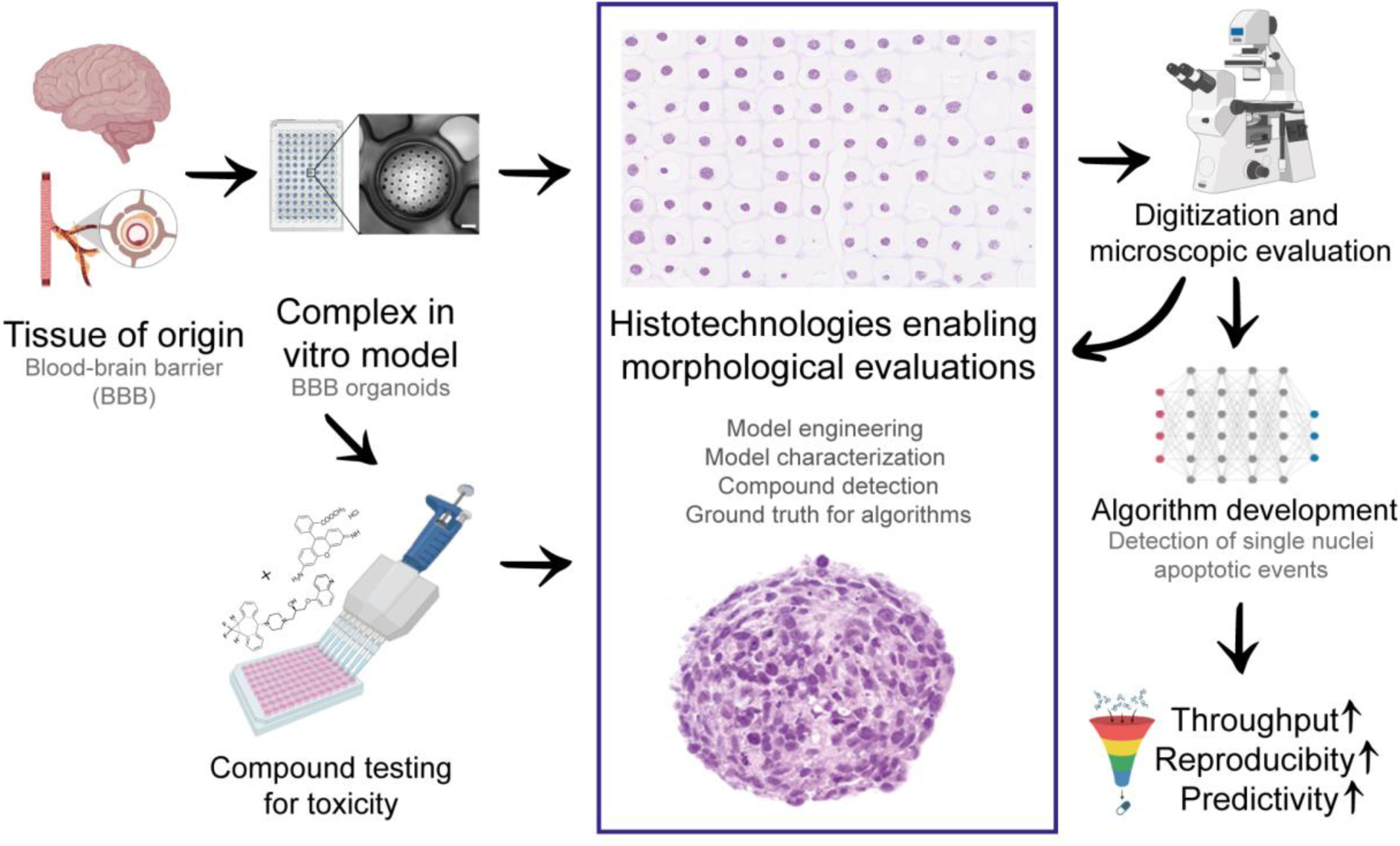

The graphical abstract was partially created with biorender.com.

**Highlights:**

- Advanced Complex *in vitro* models (CIVMs) like Blood-brain barrier (BBB) organoids show promise for preclinical drug testing. However, robust and reproducible techniques are crucial for the acceptance of CIVMs in drug development processes, especially for toxicity assessments which are highly regulated by health authorities.
- We developed orthogonal image-based readouts on histological sections to enable the use of BBB organoids for future compound toxicity assessment.
- A newly established artificial intelligence (AI) algorithm provides automated and label-free detection of apoptotic cells in drug screening using BBB organoids and provides an alternative killing assay on single cell resolution (40×) to current standards.

## Introduction

Compound attrition rates remain challenging for drug companies, with over 1 billion U.S. dollars invested per marketed drug^1^. Part of the attrition challenge is attributed to low animal-to-human translational success rates, so-called “translational failure”^2^. To overcome these shortcomings, there is a huge demand for more human predictive translational platforms and technologies, defined as New Approach Methodologies (NAMs)^3^. One promising example of NAMs are Complex *in vitro* models (CIVMs), i.e. organoids and microphysiological systems (MPS), which are complex three-dimensional (3D) “mini organ” like structures. However, the successful implementation and acceptance of CIVMs into preclinical pipelines require their proper characterization, validation and qualification to demonstrate to end-users, as well as health authorities, that they are fit for a particular purpose or context of use (CoU)^1,4,5^. Such an assessment is standard for methods used in the regulatory environment but less present in research^4,6^. At the Organization for Economic Cooperation and Development (OECD) level, there are three relevant guidance documents for *in vitro* method development and use: the Guidance Document on Good *in vitro* Method Practices (GIVIMP)^7^, the Guidance Document for Describing Non-Guideline *in vitro* Test Methods (GD211)^8^ and the Guidance Document on the validation and international acceptance of new or updated test methods for hazard assessments (GD34)^9^. GD34 and GD211 support the thorough description, characterization and validation of CIVMs/MPS in order to enhance their credibility. While characterization can include an assessment of the biological relevance and scientific validity of the CIVMs, validation refers to the process by which the reliability and relevance of a particular assay is established for a defined CoU^9,10^. Ideally, diverse teams of scientists, including engineers, biologists, and pathologists should collaboratively develop and characterize novel CIVMs, and collectively assess their CoU^11^.

To compare scientific data for characterization, qualification, and validation of novel CIVMs, the harmonization of readouts is a necessary step that allows future toxicity screening. For *in vivo* toxicity studies, histopathology is a standard readout and one of the main toxicological endpoints supporting the entry into human milestone in drug discovery. However, adapting the standard histopathology workflows from tissues to CIVMs is challenging due to their small size, differences in culturing devices, and cell composition or origin^11^. In addition, imaging and visualization of CIVMs is challenging due to factors such as microtissue thickness, light scattering, and impaired diffusion of reagents across multiple cell layers^12^. Breaking down the evaluation of these 3D models on 2D glass slides will therefore allow higher throughput by serial sectioning, while cell-cell interactions and spatial distribution are retained - similar to evaluating toxicologic pathology findings in animal organs^13^. In addition, when adapting histopathological technologies from tissues to CIVMs, recent advancements in digital pathology can be scaled-up and applied to CIVMs^11^.

A major challenge for developing drugs to treat diseases of the central nervous system (CNS) is to assess their transport and tolerability at the blood-brain barrier (BBB)^14^. While these evaluations are normally performed in animal models, there are substantial differences in the transcriptome and proteome of mice compared to humans^15^ that ultimately reduce the predictive value of rodent studies. Therefore, the recent development of CIVMs that recapitulate features of the BBB, is considered a powerful development^16^. Independent groups established a novel organoid-based model of the BBB formed by the self-organization of human brain endothelial cells, pericytes and astrocytes^17-20^. BBB organoids allow direct cell-cell interactions in the absence of artificial membranes or substrates and, importantly, recapitulate key cellular and molecular properties of the BBB, including: tight junction formation, efflux pump expression and activity and receptor-mediated transport of peptides^18^. Although this system is a major advancement in the field of *in vitro* BBB modeling, organoid production, processing, and analysis still require extensive manual work^20^. In a significant increase to process efficiency, Simonneau et al.^21^ developed BBB patterned organoid arrays based on hydrogel micro-scaffolds^22^, which allowed the formation of more than 5000 viable organoids with reproducible sizes in a 96-well plate format.

In the present manuscript, we built on the work of Simonneau et al.^21^ and Kassianidou et al.^23^ and developed new protocols by adapting standard histotechniques to enable future morphology-based toxicity assessment of drugs in BBB organoid arrays. We demonstrated the use of histotechniques to support model engineering, characterization and validation steps by using them to i) discriminate key cellular markers in BBB organoids, ii) compare those key markers to human brain tissue, iii) assess assembly and spatial distribution within the organoid using immunohistochemistry (IHC), multiplex immunofluorescence assays (mIF), and iv) localize compounds within the BBB organoids by Matrix Assisted Laser Desorption/Ionization Mass Spectrometry Imaging (MALDI-MSI). In order to automate the histopathological evaluation of toxicity, we developed an artificial intelligence (AI) algorithm that detected single-cell apoptotic events in hematoxylin & eosin (HE) 40× whole slide images (WSI) of BBB organoids on tissue microarrays (TMA).

## Results

### BBB organoids recapitulate IHC expression markers of human tissue of origin

To characterize the cellular composition and architecture of BBB organoids in relation to their human brain tissue of origin, we adapted existing paraffin embedding protocols for monoculture cells and organoids in order to account for their small size and high number of replicates. To discriminate brain endothelial cells, we chose to visualize P-glycoprotein (P-gp), one of the key efflux transporters of the BBB^24^. We used neural/glial antigen (NG)-2 and glial fibrillary acid protein (GFAP) to identify pericytes and astrocytes, respectively. Using cell pellets of monocultured cells, we detected a variable fraction of positive signal for P-gp, NG-2, and GFAP in all three cell pellets (**Supplemental Information**, **Figure 1**). Expression of P-gp, NG-2, and GFAP was detected in the human cortex (**Supplemental Information**, **Figure 2A**) and BBB organoids (**Supplemental Information**, **Figure 2B**), confirming previous data showing that the BBB organoids recapitulate the expression of markers of the BBB *in vivo*^21^.

Further characterization of the BBB organoids towards the human brain tissue of origin, using mIF with the same IHC markers provided a more detailed spatial evaluation of the different cell types within the human cortex and BBB organoids (**Figure 1**). These mIF images show that the BBB organoid is composed of an outer layer of endothelial cells and an inner meshwork of astrocytes and pericytes (**Figure 1A, B**) forming together the barrier. Ki67, which marks proliferating cells (**Figure 1C**, orange), showed equal distribution through the organoid and co-localization with all three cell components, indicating that all three cell types are viable and actively proliferating at the stage of harvesting and fixation. Caspase-3 staining (**Figure 1C**, purple) for apoptotic cells shows the baseline amount of cell death in the BBB organoid. This baseline is important for the evaluation of compound-induced cell death and is a characteristic of the BBB organoids which is not present in normal human brain tissue. Summarizing, by the selected discriminative markers and mIF technique, we can show that BBB organoids recapitulate the BBB cell composition of human brain tissue, and highlight that the barrier is at the outer edge of the organoid.

**Figure 1.**
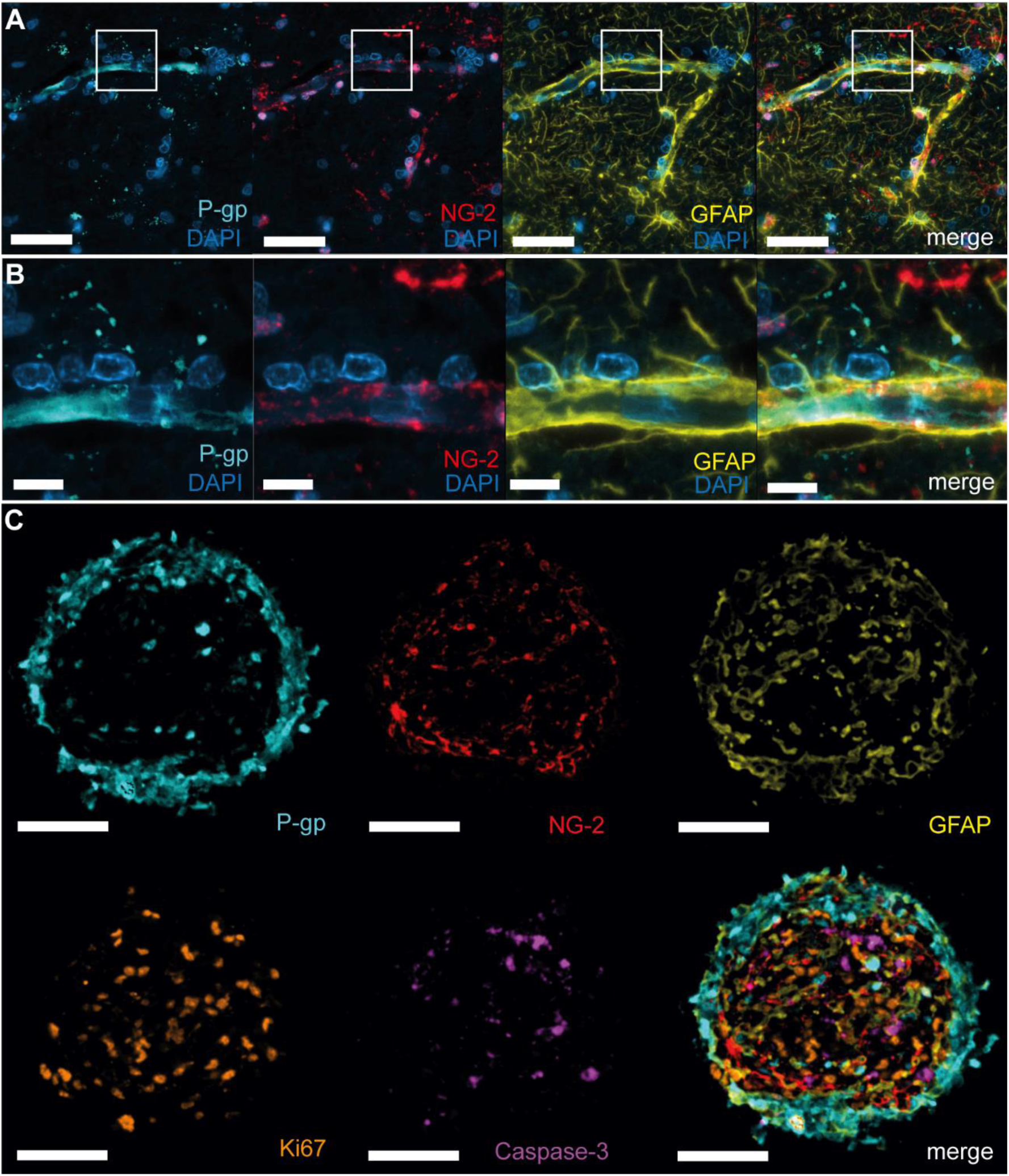
Key marker expression profile and architecture of BBB organoids compared to human brain tissue of origin. Representative images of the human cortex **(A** and **B)** and BBB organoids **(C)**. **(A** and **B)** Multiplex immunofluorescence of human cortex tissue with BBB-specific markers (P-gp for endothelial cells [turquoise], NG-2 for pericytes [red] and GFAP for astrocytes [yellow]), demonstrating similar morphologic expression patterns human brain tissue and BBB organoids **(C)**. **(C)** Co-localization and spatial distribution of cells in BBB organoid showing P-gp (turquoise) for endothelial cells, NG2 (red) for pericytes, GFAP (yellow) for astrocytes and Ki67 (orange) for proliferating cells and Caspase-3 (purple) for apoptotic cells. Scale bar, 100 μm in A, 10 μm in B, and 50 μm in C.

### Detection of rhodamine 123 as a proof of concept (POC) for the image-based readout of efflux transport in BBB organoids

Rhodamine 123 is a widely used P-gp substrate and can be used in combination with zosuquidar^25^, a highly-specific inhibitor of P-pg, to examine the functional activity of the efflux pump P-gp^26^. To functionally assess the penetration and distribution of rhodamine 123 within BBB organoids, we utilized both whole-mount widefield fluorescence microscopy and MALDI-MS imaging (**Figure 2A**). Whole mount widefield microscopy detected rhodamine 123 fluorescence at 508/528 nm, revealing its distribution within the organoids (**Figure 2B**, top and middle row). MALDI-MSI allowed the detection of rhodamine 123 at its specific mass (m/z 345.12337), when overlaying the MALDI-Fourier transform ion cyclotron resonance (FTICR) MS image onto the optical scan (**Figure 2B**, bottom row). By using this approach, a spatial map of the distribution of rhodamine 123 within the organoid section was generated. Treatment of BBB organoids with the P-gp inhibitor zosuquidar revealed a trend towards increased relative fluorescent intensity of rhodamine 123 compared to non-treated organoids, indicating a functional efflux in BBB organoids (**Figure 2C**).

**Figure 2.**
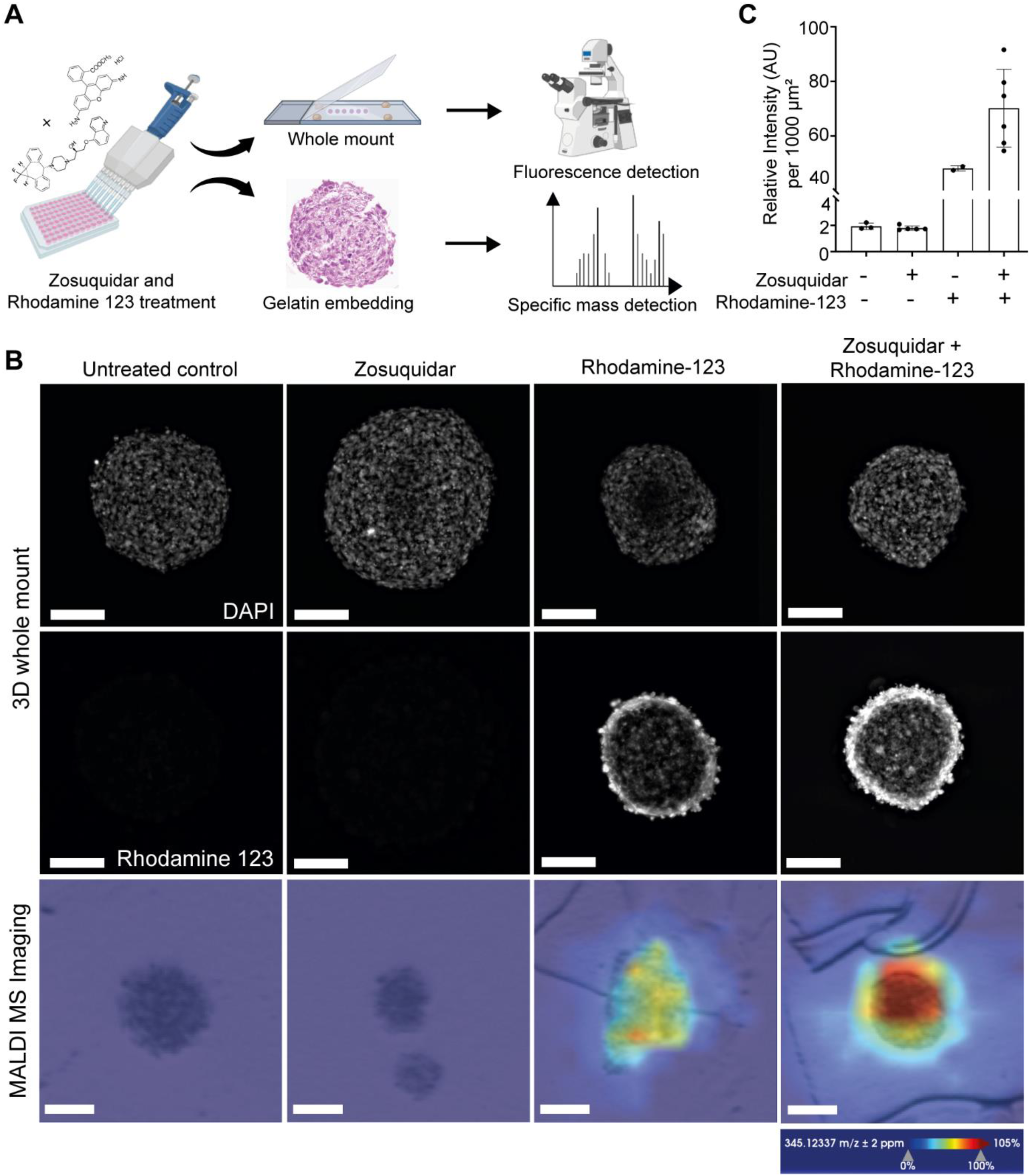
Detection of rhodamine 123 in BBB organoids by whole mount imaging and MALDI-MS imaging. **(A)** Schematic workflow of BBB organoids which were either untreated, pre-incubated with the P-gp inhibitor zosuquidar (1µM for 1 h) or the P-gp substrate rhodamine 123 (100 µM for 1 h) alone, or both in combination. Fixed BBB organoids were then embedded as whole mount on glass slides or in gelatin, subjected to either 3D whole mount fluorescence imaging or MALDI-MS imaging for specific mass detection of rhodamine 123, respectively. **(B)** Whole mount widefield microscopy imaging was performed to detect rhodamine 123 at 508/528 nm (top and middle panel). Gelatin-embedding and cryo-sections were prepared to detect rhodamine 123 at its specific mass (m/z 345.12337 Da) by MALDI-MS imaging (lower panel). Ionization mode over a mass range of 100–3000 m/z, with a lateral resolution of 50 µm mode and brightfield images are depicted. The signal for each pixel was normalized against the root mean square and scaled from 0% to 105%. Scale bars, 100 µm. **(C)** Quantification of rhodamine 123 uptake in the BBB organoid core by measuring intensity per area from whole mount imaging data. Each data point represented one whole organoid.

### High object-based accuracy for the detection of apoptotic nuclei by the newly developed automated image analysis AI algorithm on HE whole slide images of BBB organoids

To automate the histopathological evaluation of toxicity, we developed an AI algorithm that detects single-cell apoptotic events in HE 40× WSI of BBB organoid TMAs. To generate the ground truth (GT), we manually annotated normal and Caspase-3-positive apoptotic nuclei on HE slides to generate the percentage of apoptotic cells per BBB organoid (**Figure 3A**, **Supplemental Information Figure 3A**, see methods for details). The AI algorithm identified 5445 nuclei, slightly surpassing the 5422 nuclei originating from the five organoid sections previously annotated by pathologists. Additionally, it identified 3293 apoptotic nuclei, while pathologists have previously annotated 2250 apoptotic nuclei from 39 organoids that had been used for algorithm training. To assess the accuracy of the AI algorithm, we employed both pixel-based and object-based accuracy metrics (**Supplemental Information Figure 3B** and **C**); see representative images of nuclei and apoptotic detection (**Supplemental Information**, **Figure 4**). The AI algorithm’s performance was quantified using sensitivity, specificity, and F1 scores. These metrics provide a comprehensive evaluation of the AI algorithm’s ability to detect true positives, true negatives, false positives, and false negatives. The high sensitivity and recall values indicate that the AI algorithm is effective in identifying apoptotic nuclei, while the precision values highlight its accuracy in minimizing false positives (**Table 1**).

**Figure 3.**
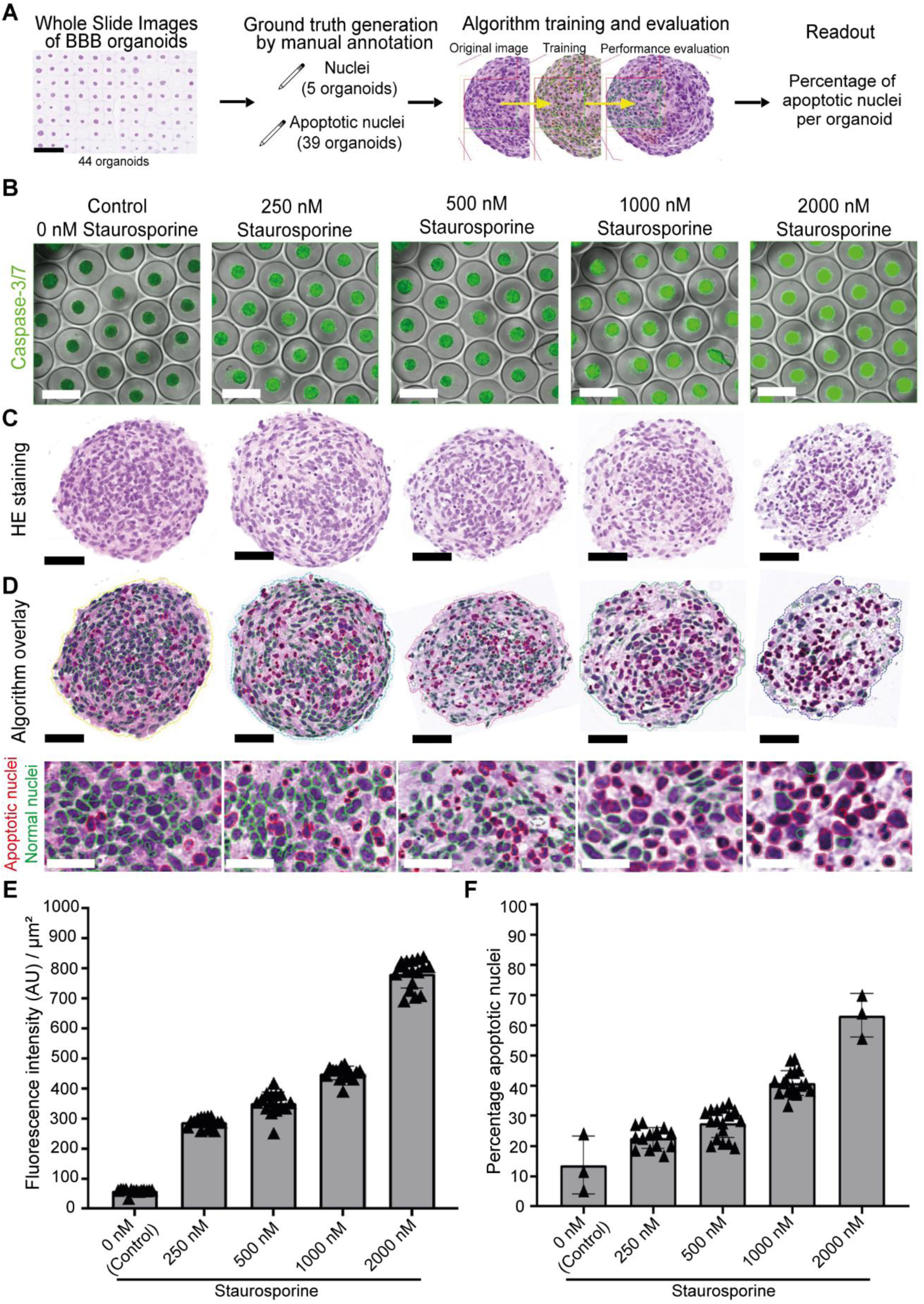
Detection of staurosporine-induced apoptosis in BBB organoids by the standard image-based Caspase-3/7 assay and the newly developed automated image analysis AI algorithm in comparison. **(A)** Overview of the workflow for the AI algorithm development. HE WSI of BBB organoids were used to generate GT by manual annotation of nuclei and apoptotic nuclei. Annotation labels on HE slides went into the AI algorithm training. The readout of the AI algorithm is the percentage of apoptotic cells per BBB organoid. **(B)** Representative fluorescent images of Caspase-3/7 (green) signal overlaid on brightfield images of BBB organoids, treated with different concentrations of staurosporine for eight hours. Caspase-3/7 CellEvent was added for 30 min before live imaging at 5×. Scale bar, 500 µm. **(C)** Representative HE images of BBB organoids, scanned at 40×. Scale bar, 50 µm. **(D)** Algorithm overlay showing the detection of apoptotic (red outline) and normal nuclei (green outline) by the newly developed AI algorithm and magnified inserts of the same organoids treated with different concentrations of staurosporine for eight hours. Scale bar in upper panel, 50 μm, scale bar of inserts in lower panel, 20 µm. **E -F**, Quantification of apoptotic events in BBB organoids treated with different concentrations of staurosporine. **(E)** For the standard image-based Caspase-3/7 assay, mean fluorescent intensity per area was quantified. Each data point represented one whole organoid. Mean ± SD are depicted. **(F)** As detected by the newly developed AI algorithm, the percentage of apoptotic nuclei of total nuclei per organoid section was quantified. Each data point represented one organoid section. Mean ± SD are depicted.

**Table 1.** Scoring of the newly developed automated image analysis AI algorithm to detect apoptosis in BBB organoids.

| Data set | Accuracy | Ground truth | Precision | Recall | F1 Score* |
| --- | --- | --- | --- | --- | --- |
| Training organoids | Pixel-based | Nuclei segmentation | 0.978 | 0.982 | 0.980 |
| Training organoids | Pixel-based | IHC Caspase-3 | 0.341 | 0.770 | 0.460 |
| Training organoids | Object-based | Pathologist annotation | 0.767 | 0.884 | 0.817 |
| Naive organoids | Object-based | Pathologist annotation | 0.478 | 0.709 | 0.571 |

\*F1 Scores above 0.7 are considered high accuracy, whereas scores around 0.5 indicate moderate accuracy and <0.5 are considered low accuracy.

Pixel-based accuracy was determined by calculating the overlap between the training labels and the AI algorithm’s output labels. Pixel-based accuracy for nuclei segmentation on training organoids was almost perfect (precision, P = 0.978, recall, R = 0.982, and F1 score = 0.98), while for the Caspase-3 IHC precision was rather low (P = 0.341), but had much higher recall (R = 0.770), resulting in a moderate F1 score (F1 = 0.460). When compared to the object-based accuracy based on the pathologists’ annotations, we observed an increase in higher precision, recall and F1 score (P = 0.767; R = 0.884; F1 = 0.817). Of note, the agreement among three blinded European board-certified veterinary anatomical pathologists was also evaluated to ensure the robustness of the AI algorithm. The high inter-rater accuracy of M = 0.93 and reliability of ĸ = 0.85 underscores the reliability of the AI algorithm in replicating expert annotations. Object-based accuracy was additionally evaluated on naive organoids that were treated with different concentrations of staurosporine for eight hours (see methods for details), or left untreated. Here, the recall was 0.709, precision was 0.478, and the F1 score was 0.571. These metrics indicate that the AI algorithm also performs well on previously unseen data.

### The newly developed automated image analysis AI algorithm compares favorably to the standard image-based 5× Caspase-3/7 assay in detecting apoptosis in BBB organoids

Performance of the newly developed AI algorithm was validated against the standard image-based Caspase-3/7 assay that contains fluorogenic substrates for activated caspase-3/7 and therefore allows detection of apoptosis in live cells. To this end, we induced apoptosis dose-dependently using staurosporine in BBB organoids as a benchmark to generate BBB organoids which can be used to quantify apoptosis as a future toxicity readout (**Figure 3B-F**)^27^. For the standard image-based Caspase-3/7 assay at 5× magnification, fluorescent intensity per area was measured (**Figure 3B** and **E**). In parallel, the AI algorithm was applied to a HE WSI of a TMA of BBB organoids quantifying the percentage of apoptotic nuclei relative to the total nuclei per organoid section at 40× magnification (**Figure 3C, D**). Both assays capture the staurosporine induced, dose-dependent increase in apoptosis over time (**Figure 3E, F).** Comparing both readouts, the AI algorithm provides an equally robust and reliable method of apoptosis detection as compared to the current standard image-based Caspase-3/7 assay and is therefore a suitable tool to assess drug-induced apoptosis. In addition, the AI algorithm is superior in detecting low frequency apoptotic events, as indicated by the higher baseline in the untreated organoids (**Figure 3F)**, similar to the microscopic evaluation by a pathologist.

## Discussion

Novel technologies around CIVMs biology are evolving. Nevertheless, there is a lack of simple, standardized and high throughput technologies to evaluate microtissue (i.e organoid) morphology and functionality. Morphologic assessment of CIVMs can provide key information for the characterization and validation of a novel model including the identification of cell-types present and features of cellular viability and maturity compared to normal tissues^11^. By using BBB organoids, we showed that histotechniques are simple, robust and highly standardized methodologies which can be transferred from regulatory toxicologic pathology assessments *in vivo* to *in vitro*. Importantly, the histotechniques presented here allow the direct comparison and quantification of IHC signals within the human brain tissue of origin and the BBB organoids, using identical antibodies and staining protocols. Another advantage of adapting standard histotechniques to CIVMs in the regulatory environment is that regulators are familiar with the interpretation of morphologic evaluations performed by pathologists.

With the adapted histotechnology data (IHC & mIF), we were able to reproduce previous BBB model characterizations by fluorescence microscopy^17,18,20,21^, and, importantly, achieve an increase in throughput, automation, and standardization. Specifically, combining BBB organoids from different treatment groups into one TMA^28^ offers a high-throughput approach for processing and evaluation. This enables the user to perform routine HE, special stains, and IHC on multiple samples in one paraffin block, removing slide-to-slide and run-to-run variability and allowing for automated and AI based quantitative image analysis, such as those shown here. In addition, the automated sequential staining and multispectral imaging allowed for precise spectral unmixing and high-resolution visualization of the cellular components and co-localization of different functional (e.g. P-gp, Caspase3, Ki67) and morphological markers (NG-2, GFAP).

As a functional barrier, the BBB fulfills its primary function of protecting the brain by controlling the access of solutes and toxic substances to the CNS. Assessment of compound penetrance of the BBB is a standard assay along the drug development value chain. Analysis of high molecular weight biomolecules with high detection accuracy and sensitivity can be achieved by MALDI-MS^25,29,30^, rendering it a suitable method to assess compound penetration in organoids. As a POC to develop MALDI-MSI on organoids, we assessed rhodamine 123 penetration^26^ in BBB organoids treated with zosuquidar, a highly specific P-gp inhibitor^25^. Along with the experiment, we established image-based functional readouts with MALDI-MS and widefield microscopy to demonstrate morphologically the retention of rhodamine 123 in BBB organoids upon P-gp inhibition by Zosuquidar. While widefield microscopy requires compounds that are inherently fluorescent or are labeled with a fluorophore^20^, MALDI-MSI is a reliable method that can assess BBB penetrance of novel compounds in development (especially small molecules), where fluorophore labelling is not applicable^31^. For compound distribution where single cell resolution is required, light and fluorescence microscopy methods, e.g. IHC for large molecules, *in situ* hybridization (ISH) for Antisense Oligonucleotides (ASOs) and Gene Therapy modalities may be more suitable and should be chosen over MALDI-MS.

The current gold standard for toxicologic pathology (morphologic) evaluation is the manual evaluation, scoring and interpretation of HE tissue sections by light microscopy^13,30^. To transfer this workflow to CIVMs is for a pathologist cumbersome, too costly and time consuming^32^. In addition, CIVMs have a baseline of “normal” apoptotic and proliferative cells (as shown by the characterization data by mIF), which makes quantitative image analysis necessary. Therefore, the presented newly developed AI algorithm provides an automated solution to perform quantitative morphologic readouts at the single cell level in BBB organoids. In addition, the results corroborated the findings from the standard image-based Caspase-3/7 assay at 5× magnification, demonstrating the robustness and sensitivity of the newly developed AI algorithm in detecting apoptosis in 40× HE WSI TMA digital scans. Interestingly, the AI algorithm applied on standard HE sections outperforms the performance obtained from the Caspase-3 IHC staining, which is reflected in the increased F1 score in pathologist’ annotations. This suggests that the AI algorithm detects histomorphologic features and patterns beyond the Caspase-3 staining, similar to manual evaluations by pathologists, and is therefore superior to the standard quantitative image analysis of a Caspase-3 IHC.

Compared to 3D imaging approaches, 2D analyses, like the advanced histotechnologies demonstrated here, have significant benefits in terms of computational efficiency, algorithmic complexity and ease of data handling and interpretation. However, certain spatial relationships and interactions can only be accurately assessed in 3D. In a first POC study, we found similar results for the percentage of apoptotic nuclei in 2D HE scans and 3D reconstructed DRAQ5 BBB organoids that have been treated with different concentrations of staurosporine (data not shown). This warrants further validation experiments, but our initial data suggests that 2D is superior in throughput and reproducibility, while requiring significantly less computational power and time to process the images. We therefore recommend the preferential establishment of 2D imaging applications, like the advanced histotechnologies for morphologic evaluations of e.g. toxicity screening assays with CIVMs.

In conclusion, advanced tissue technologies (HE, IHC, mIF, MALDI-MS) can be adapted to CIVMs such as BBB organoids. They are examples of simple, robust and highly standardized methodologies which can be transferred from regulatory toxicologic pathology assessments *in vivo* to *in vitro*. The presented POC data with tool compounds demonstrate the potential use and benefits of integrating advanced histomorphologic-based readouts and digital pathology AI algorithms on CIVMs for model development, characterization and validation for specific CoU.

With this interdisciplinary approach of combining novel CIVMs with highly standardized and regulated methodologies, we aim to improve the throughput, reproducibility and predictivity of preclinical drug development in the future when using CIVMs. In addition, we foster the interdisciplinary exchange and data comparison along the drug development value chain. As a consequence, we aim to enable the future acceptance of NAMs for benefit-risk assessment. We encourage the scientific community to explore similar options for their projects and to engage with health authorities on the use of morphologic readouts for CIVMs.

### Limitations of this study

IHC marker expression in cell pellets of origin can differ from that of assembled organoids, due to factors like number of cell passages, media used, co-culture or impact by extracellular matrices. IHC is not sensitive enough to detect low protein expression levels and does not provide a quantitative method to assess expression levels of the selected discriminative markers in BBB organoids. MALDI-MSI faces throughput limitations, with slide-by-slide acquisition balancing resolution (here 50 µm) and sensitivity, defining the limit of detection, therefore often yielding qualitative rather than quantitative data. In the context of AI algorithm evaluation, ground truth validation by pathologists is crucial, as they must become familiar with CIVM histology, including baseline apoptotic events in static cultures, to ensure inter-rater and intra-rater reliability, necessitating specific training. However, pathologist evaluations are also subjective to a certain extent. We therefore included an objective ground truth using the Caspase-3 staining, but it is important to keep in mind that the current AI algorithm, trained on histomorphological features, is somewhat general and may not differentiate between apoptosis, necrosis, or other cell death mechanisms.

## Experimental Procedures

### Key resource table

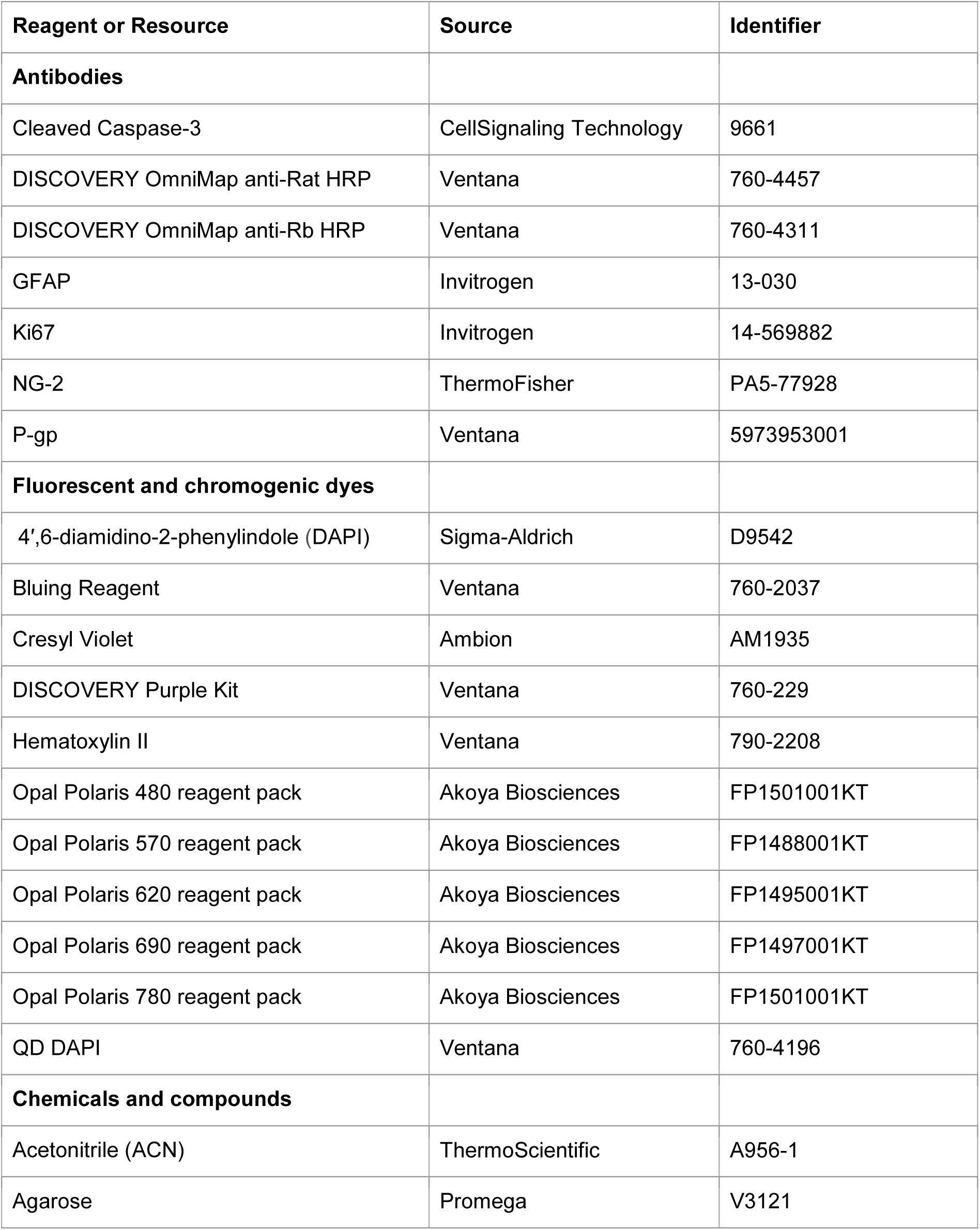
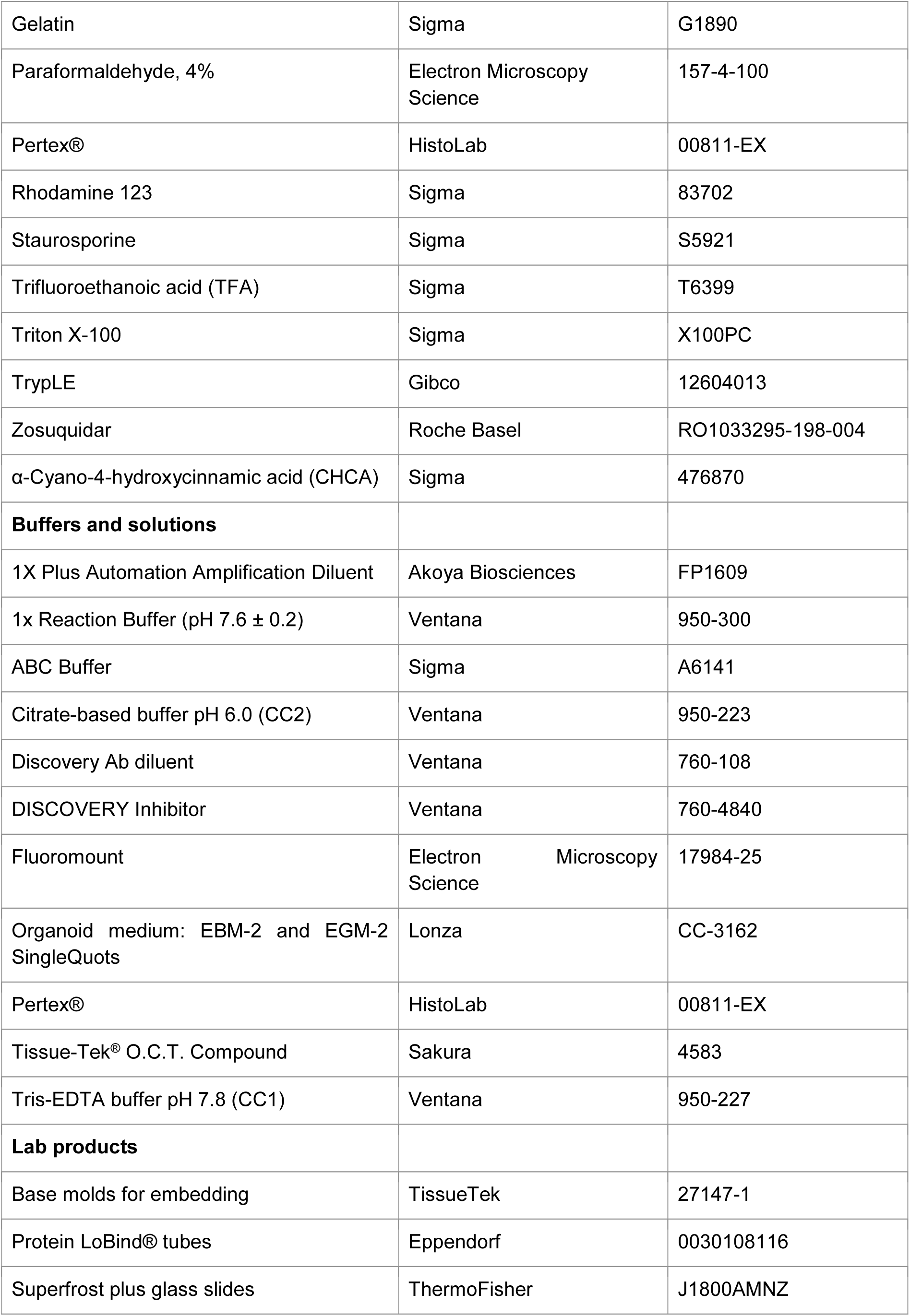
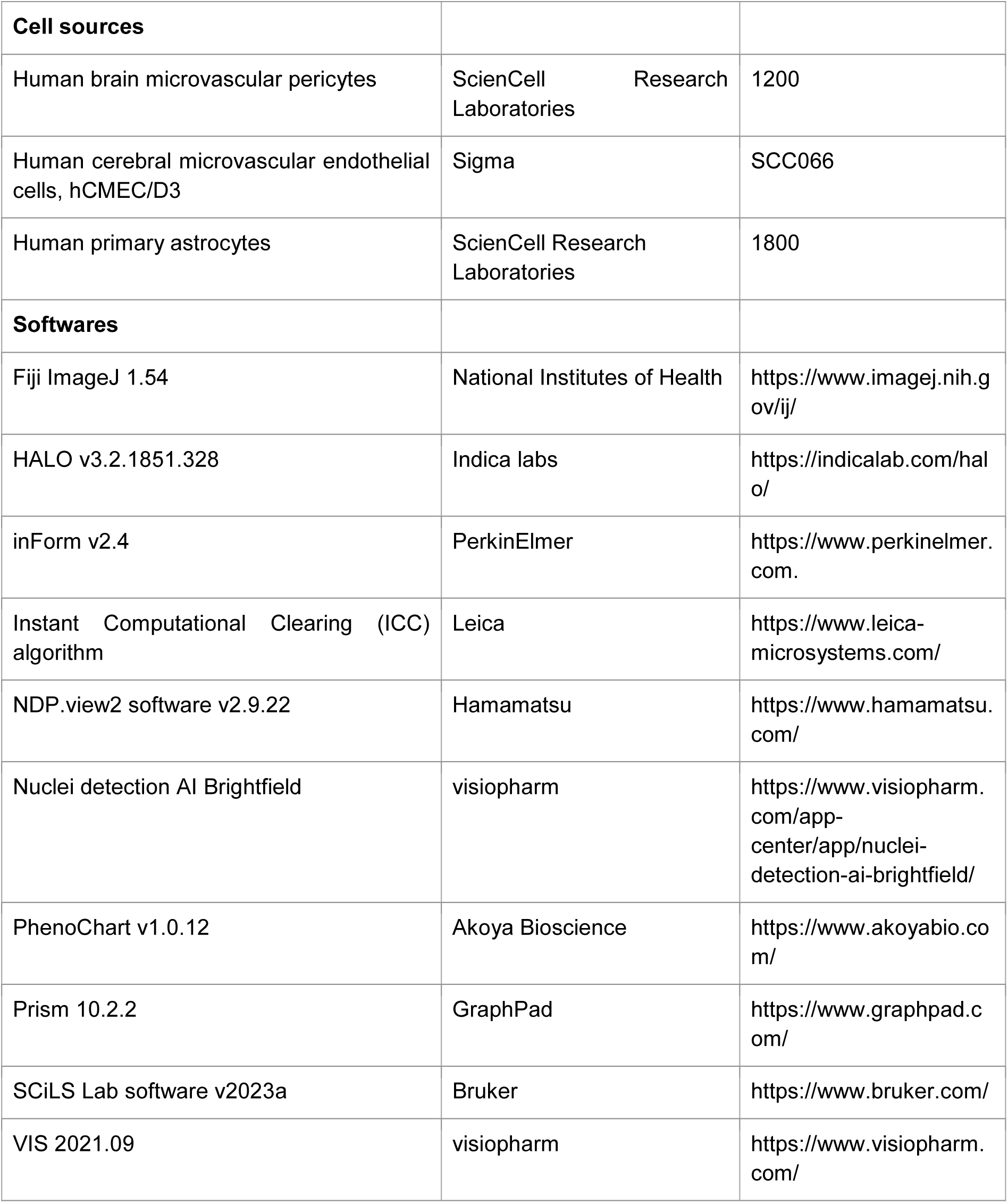

### Lead contact and materials availability

All unique/stable reagents generated in this study are available from the Lead Contact, Nadine Stokar-Regenscheit, without restriction.

### Data and code availability

The data that was used as GT for the algorithm development are available from the corresponding author (N.S.-R.), upon reasonable request.

### Generation of BBB organoids and information on cell sources

The creation of BBB organoids has been previously described by Simonneau et al.^21^. In the present study, individual cell lines and BBB organoids were generated following the protocol by Kassianidou et al.^23^. Cell line sources are indicated in the key resource table.

### Human brain tissue sample source

For validation of key cellular markers by multiplex mIF, one formalin fixed and paraffin embedded (FFPE) block from the frontal cortex of a healthy male Caucasian donor (83y) was purchased from Asterand Bioscience. All human tissue samples supplied by ASTERAND® At BioIVT, were collected under IRB-approved protocols and are de-identified prior to banking and distribution. BioIVT complies with the US Health Insurance Portability and Accountability Act (HIPAA), the United Kingdom Human Tissue Authority (HTA) and is accredited by the College of American Pathologists (CAP).

### PFA fixation and paraffin embedding of BBB organoids and cell pellets

After successful self-assembly, BBB organoids were fixed using 4% paraformaldehyde (PFA, 157-4-100, Electron Microscopy Science) for 20 minutes at room temperature (RT), followed by three washing steps in phosphate buffer saline (PBS). BBB organoids from several wells were collected in a Protein LoBind® tube (0030108116, Eppendorf) by flushing them out with 0.1% Triton-X (X100PC, Sigma) in PBS and were allowed to settle before removing the supernatant. Organoids were visualized via staining with 10 µL eosin for one minute (150 mL eosin, 15 mL phloxine, 4 mL acetic acid, 780 mL ethanol 96%) and washed three times in PBS to remove excessive eosin. The supernatant was then carefully removed using a pipette tip attached to an aspirator to avoid aspirating the organoids, while leaving approximately 50 µL in the tube. Agarose (V3121, Promega) was diluted to 2% in distilled water, boiled until completely dissolved, and kept at 56 °C until use. The organoids were transferred to disposable base molds (27147-1, TissueTek), and then 250 µL of liquid agarose was added on top of the organoids. Immediately, the molds were centrifuged at 200 rpm for 2 minutes at RT to ensure all organoids settled in a similar plane. Molds were then placed on ice to allow the agarose to solidify. Several inserts of molds were collected in one biopsy cassette (CXN3.1, Carl Roth) and dehydrated overnight using a vacuum filter processor (TissueTek VIP5, Sakura, protocol: 2x 70% ethanol, 30 minutes, 35°C; 1x 80% ethanol, 60 minutes, 35°C; 1x 96% ethanol, 30 minutes, 35°C; 2x 100% ethanol, 60 minutes, 35°C; 2x xylol 45 minutes, 35°C; 4x paraffin, 45 minutes, 60°C). Using a tissue embedding console (Tissue-Tek® TEC™5, Sakura), we embedded up to eight inserts of dehydrated organoids in liquid paraffin with the organoids facing down in the metallic mold. After letting the paraffin blocks dry on ice for 30 minutes, blocks were stored at RT until cutting with a microtome. For the generation of cell pellets, adherent cells (primary human astrocytes, ScienCell Research Laboratories; Human brain microvascular pericytes, ScienCell Research Laboratories; Human cerebral microvascular endothelial cells, hCMEC/D3, Sigma Aldrich, see Simonneau *et al.* ^21^ for details) were detached using TrypLE (12604013, Gibco), collected in their respective culture medium and centrifuged at 1500 rpm for 5 minutes at RT. After removing the supernatant, cell pellets were washed in PBS once and centrifuged again with the same settings. After aspirating PBS supernatent, cells were immediately fixed in 4% PFA for 30 minutes, followed by three washes in PBS. The same protocol as BBB organoids described above was applied for PFA-fixed paraffin embedding of cell pellets.

### Rhodamine 123 treatment for MALDI-MS detection

Multicellular BBB organoid arrays were formed for 48 h and consequently pre-incubated with zosuquidar (RO1033295-198-004, Roche Basel, 1 µM) in Endothelial Basal Medium (EBM-2, Lonza) supplemented with hydrocortisone, GA-1000, 5% FBS, hEGF, hFGF-B, R3-IGF-1, ascorbic acid and heparin (EGM-2 SingleQuots Supplements, Lonza) medium without VEGF and reduced FBS (2%) for one hour at 37°C with 5% CO_2_. The samples were then treated with rhodamine 123 (83702, Sigma, 100 µM) for one hour with or without zosuquidar to assess active uptake. After treatment, BBB organoids were washed three times with 37°C warm medium and then fixed in 4% PFA for 20 minutes at RT. Samples were washed thoroughly again, transferred to Protein LoBind® tubes (0030108116, Eppendorf) and embedded in gelatin for MALDI-MS detection as described below.

### Gelatin embedding and sectioning of BBB organoids for MALDI-MS detection

BBB organoids were gelatin-embedded for MALDI-MSI using a method adapted from Bakker et al. (2022). Briefly, BBB organoids from several wells of GRI3D 96-well plate (MSPPGRI3D-96P-S) were collected in a Protein LoBind® tube (0030108116, Eppendorf) by flushing them out with 0.1% Triton X-100 (X100PC, Sigma) in PBS and allowed to settle before removing the supernatant. To allow visualization, organoids were stained with cresyl violet (AM1935, Ambion) for 30 seconds, followed by three PBS washes. ABC buffer (50 mM NH4CO3, A6141, Sigma) was subsequently added, and organoids were resuspended in 37°C 15% gelatin solution (G1890, Sigma), and immediately transferred to a disposable base mold (27147-1, TissueTek) before being covered with gelatin and rapidly frozen in liquid nitrogen. Blocks were kept at -80°C until cutting with a cryostat. Frozen sections were cut at 10 µm and mounted on Superfrost plus glass slides (J1800AMNZ, ThermoFisher).

### Whole-mount immunofluorescence staining and widefield microscopy

For imaging purposes, rhodamine 123 and zosuquidar treated BBB organoids were flushed from the plate with PBS with 0.1% Triton-X (X100PC, Sigma) and transferred into Protein LoBind® tubes (0030108116, Eppendorf) and allowed to settle for 2-3 minutes before removing the supernatant. 4′,6-diamidino-2-phenylindole (DAPI) (D9542, 1 μg/mL, Sigma) was then added to the tubes and incubated for one hour at RT on a rotary shaker in the dark. Finally, the samples were washed again with PBS, transferred to cover glasses, and mounted with Fluoromount (17984-25, Electron Microscopy Science). Organoids were imaged using a Leica Microsystems, Thunder Imager 3D Assay with a 20 × / 0.55 Ph2 dry objective. The images were acquired with a 2x2 binning in a 16 bit format. Two different channels were set up; one for DAPI (UV laser), and one that matches the fluorophore of the rhodamine 123, in this case Alexa Fluor 488 (488 laser). A z-stack covering a total depth of 8.5 μm, using 8 steps with the core placed at the center (1.21 µm step size) was used. The Instant Computational Clearing (ICC) algorithm by Leica was then applied to the images.

### TMA generation of BBB organoids

PFA fixed BBB organoids were embedded into a TMA by Micromatrices as described in Plummer *et al.* ^28^.

### Histology of PFA fixed and paraffin embedded (PFA-PE) organoids

From each PFA fixed and paraffin embedded (PFA-PE) block, 4 µm thick sections were cut and mounted on Superfrost plus glass slides (J1800AMNZ, ThermoFisher). Slides were incubated in a slide oven overnight at 37°C with the specimen facing up to prevent loss of the small organoids due to the melting paraffin. HE staining was performed (Ventana HE600, Roche) according to the manufacturer’s standard protocol.

### Automated immunohistochemistry PFA-PE sections

IHC on PFA-PE slides was performed using Ventana Discovery Ultra automated tissue stainer (Roche Tissue Diagnostics, Tucson AZ USA). Slides were baked at 60°C for eight minutes and subsequently warmed further to 69°C for 24 minutes, for deparaffinization. Heat-induced antigen retrieval was performed with Tris-EDTA buffer pH 7.8 (CC1, 950-227, Ventana, for Ki67 and Caspase-3) or citrate-based buffer pH 6.0 (CC2, 950-223, Ventana, for all other antibodies) at 95°C for 32 minutes (**Table 1**). After blocking with DISCOVERY Inhibitor (760-4840, Ventana) for 16 minutes, primary antibodies diluted in Discovery Ab diluent (760-108, Ventana) were added as shown in **Table 1**. Anti-species secondary antibodies conjugated to horseradish peroxidase (HRP) (DISCOVERY OmniMap anti-Rb HRP, 760-4311, Ventana; or DISCOVERYOmniMap anti-Rt HRP, 760-4457, Ventana respectively) were applied for 16 minutes and subsequent visualization was performed using DISCOVERY Purple Kit (760-229, Ventana). Lastly, specimens were counterstained with hematoxylin (Hematoxylin II, 790-2208, Ventana; Bluing Reagent, 760-2037, Ventana) for eight and four minutes, respectively. After dehydration and xylol baths, slides were aqueous mounted (Pertex®, 00811-EX, HistoLab) and dried overnight.

### Automated multiplex immunofluorescence on PFA-PE sections

PFA-PE slides were subjected to mIF using Ventana Discovery Ultra automated tissue stainer (Roche Tissue Diagnostics, Tucson AZ USA). Slides were baked at 60°C for eight minutes and further warmed to 69°C for 24 minutes for deparaffinization. Heat-induced antigen retrieval was performed with citrate-based buffer pH 6.0 (CC2, 950-223, Ventana for all antibodies) at 95°C for 32 minutes.

Slides were incubated with primary antibodies (**Table 2**) diluted in 1X Plus Automation Amplification Diluent (FP1609, Akoya Biosciences) in the following sequential order: P-gp-Opal480, NG2-Opal570, Ki67-Opal620, Caspase-3-Opal690, GFAP-Opal780. Primary antibodies were detected using anti-species secondary antibodies conjugated to HRP (DISCOVERY OmniMap anti-Rb HRP, 760-4311, Ventana; DISCOVERY OmniMap anti-Rat HRP, 760-4457, Ventana). For each sequence, the respective Opal dye (Opal Polaris 480 reagent pack, FP1501001KT, Akoya Biosciences; Opal Polaris 570 reagent pack, FP1488001KT, Akoya Biosciences; Opal Polaris 620 reagent pack, FP1495001KT, Akoya Biosciences; Opal Polaris 690 reagent pack, FP1497001KT, Akoya Biosciences; Opal Polaris 780 reagent pack, FP1501001KT, Akoya Biosciences) was applied. Slides were mounted with ProLong Glass Antifade Mountant (P36980, ThermoFisher) and dried overnight. Neutralization of HRP and denaturation of the proteins was performed after every primary antibody sequence in order to avoid cross-bleeding and cross-reacting antibodies.

**Table 2.** Primary and secondary antibodies used for immunohistochemistry and multiplex immunofluorescence stainings.

| Antibody | Vendor | Identifier | Species | Blocking<br>[y/n] | Dilution | Incubation<br>time<br>[min] | Incubation<br>Temperature<br>[°C] | Secondary<br>Antibody<br>for mIF | Target cell<br>types |
| --- | --- | --- | --- | --- | --- | --- | --- | --- | --- |
| P-gp | Ventana | 5973953001 | rb | y | Pre-diluted | 60 | 37 | Opal480 | Brain<br>endothelial<br>cells |
| NG2 | Thermo<br>Fisher | PA5-77928 | rb | y | 1:150 | 60 | 37 | Opal570 | Pericytes |
| Ki67 | Invitrogen | 14-569882 | rt | y | 1:500 | 60 | RT | Opal620 | Proliferating<br>cells |
| Cleaved<br>Caspase-3 | Cell<br>Signaling<br>Technology | 9661 | rb | y | 1:100 | 60 | 37 | Opal690 | Apoptotic<br>cells |
| GFAP | Invitrogen | 13-030 | rt | n | 1:200 | 60 | 37 | Opal780 | Astrocytes |

rb, rabbit; rt, rat; y, yes; n, no; RT, room temperature.

For validation of the key human brain cellular markers, primary antibodies targeting P-gp, GFAP and NG2 (**Table 2**) were diluted in Discovery Ab diluent (760-108, Ventana) and subjected to the same procedure as described above. Primary antibodies were added in a sequential order as follows: P-gp-Opal480, GFAP-Opal570, NG2-Opal690, counterstained with QD DAPI (760-4196, Ventana) for eight minutes.

### Chemical destain and re-stain method for Caspase3 IHC to generate GT for algorithm development

The Caspase-3 IHC staining (as described above) was applied on the same HE sections, using a chemical destain and IHC restain protocol adapted from Hinton et al ^33^. Briefly, after HE scanning, slides were heated up at 100°C to gently remove the coverslip, before the specimen was exposed to xylene, ethanol and 1x Reaction Buffer (pH 7.6 ± 0.2, 950-300, Ventana), which chemically destained the HE staining. Following this procedure, IHC for Caspase-3 was immediately performed (see methods section for Automated immunohistochemistry PFA-PE sections and **Table 1** for details), slides were mounted and scanned again.

### Whole slide scanning of HE, IHC and mIF

HE and IHC slides were imaged with a whole slide scanner at 40× (Hamamatsu, NanoZoomer S360, standard HE and IHC imaging protocol). Images were saved in .ndpi format and viewed with the NDP.view2 software (v2.9.22). Acquisition resolution was 0.23 µm/px in all images, corresponding to 40× magnification. Multiplex IF stainings using the Opal dyes from Akoya were digitized with multispectral imaging by the Vectra® Polaris™ (PerkinElmer) using the MOTiF™ technology at 40× magnification for all 5 fluorochromes. The colors were unmixed with PhenoChart (v1.0.12) and inForm (v2.4). Raw image data was saved in .qptiff, and tiles were fused in HALO (Indica labs, v3.2.1851.328). Pixel size was 0.50 µm/px in those images.

### MALDI-MS imaging

For matrix coating of BBB organoid sections on Superfrost glass slides a solution of α-cyano-4-hydroxycinnamic acid (CHCA, 476870, Sigma) at a concentration of 10 mg/mL dissolved in acetonitrile (ACN, A956-1, ThermoFisher):0.1% 2,2,2-trifluoroethanoic acid (TFA, T6399, Sigma) in water (1:1) was applied using a HTX M5 sprayer (HTX Technologies, Chapel Hill, NC). Four spraying cycles were applied at a flow rate of 0.4 mL/minute using a nozzle temperature of 90°C, a nozzle velocity of 3400 mm/minute, and track spacing of 3 mm (HH pattern). Tissue samples were analyzed on a 7 Tesla SolariX XR FTICR MS instrument with a MALDI source equipped with a Smartbeam-II^TM^ Laser System (Bruker Daltonik GmbH, Bremen, Germany). Ions were detected in positive ionization mode over a mass range of 100–3000 m/z, with a lateral resolution of 50 µm. 250 laser shots were applied per pixel at a frequency of 1000 Hz, with a laser power of 25%. The signal for each pixel was normalized against the root mean square of all data points. The ion image was co-registered with an optical scan generated from the sample. All MS images were generated with SCiLS Lab software (Version 2023a Core, Bruker). Rhodamine 123 was identified by matching measured m/z values with the value calculated based on its elemental formula (C_21_H_16_N_2_O_3_), within a mass window of 2 ppm.

### Staurosporine treatment of BBB organoids

BBB organoids were treated with different concentrations (250 nM; 500 nM; 1000 nM; 2000 nM) of staurosporine (S5921, Sigma) for eight hours at 37°C 5% CO_2_ and subsequently fixed with 4% PFA for 20 min at RT. An untreated BBB organoid group served as a negative control. For each concentration, including the negative control, five wells with approx. 55 organoids per well were used for the AI algorithm validation study.

### Automated image analysis AI algorithm development

#### Workflow

The image analysis was performed using the proprietary software VIS (version 2021.09, Visiopharm, Denmark). The morphology-based algorithm was developed to detect single-nuclei apoptosis events in HE 40× WSI of BBB organoids. To that end, BBB organoids were generated as described previously ^21,23^. BBB organoids were embedded as PFA-PE organoid pellets and cut at 3 µm, as described above. HE staining was performed according to the manufacturer’s standard protocol, while Caspase-3 IHC was performed (prepared by the chemical destain and re-stain method described above) WSI at 40× magnification was used to digitize the slides. For the AI algorithm development, 44 organoids from 14 WSIs were used for GT generation, followed by AI algorithm training and evaluation (Supplemental Information **Figure 3A**).

#### AI algorithm design and optimization

The AI algorithm was designed in a two-step approach using machine learning. First, 5422 manual annotated nuclei coming from 5 slices of unique BBB organoids from 3 different HE slides (classes: nuclei, cytoplasm, background) were fed in a pre-trained U-Net classifier (approximal 235000 iterations, #10167, Visiopharm, https://visiopharm.com/app-center/app/nuclei-detection-ai-brightfield/). In a second step, additional 2250 apoptotic nuclei labels from 39 organoids from 11 WSI (22 untreated organoids, 17 treated with 500 nM of Staurosporine for eight hours as described above) fed into the previously trained AI algorithm (47644 additional iterations). The training labels were obtained by manual annotation of nuclei in HE slides that previously showed a Caspase-3 IHC signal. Post-processing steps of the AI algorithm included a minimum size of 6 μm^2^ per nuclei and 3 μm^2^ per apoptotic nuclei. To test the performance of the AI algorithm, we used two types of GT data for the nuclei segmentation and detection of apoptotic nuclei in the HE scans: First, objective GT by Caspase-3 IHC, which was used to generate training set, and second, subjective GT by pathologists annotation (three European board-certified veterinary anatomical pathologists).

#### AI algorithm validation

AI Algorithm validation consisted of 4 steps:

1. <u>Validation against objective ground truth using Caspase-3 IHC</u>

Using the Visiopharm Software, objective GT was generated by aligning Caspase-3^+^ IHC WSI and HE WSI (**Supplemental Information Figure 3A)**. Subsequent annotations of apoptotic nuclei, identified by Caspase-3 IHC in HE WSI slides were used for training of the AI algorithm. For evaluation of the performance of the AI algorithm, pixel-based accuracy was assessed by calculating the overlap of the training labels inside the AI algorithm output labels (**Supplemental Information Figure 3B)**. This resulted in a classification as true positive, true negative, false positive, and false negative areas, respectively. Sensitivity, specificity and F1 scores were calculated accordingly.

2. <u>Validation against subjective ground truth using pathologist’s annotation</u>

The subjective GT included evaluation and manual annotation of apoptotic nuclei on HE slides by three blinded European board-certified veterinary anatomical pathologists on the same 44 BBB organoids. Object-based accuracy was assessed by comparing the nuclei classification (normal *vs.* apoptotic) pathologist’s annotation to the AI algorithm output. This resulted in true positive, true negative, false positive, and false negative nuclei, respectively (**Supplemental Information Figure 3C**). Sensitivity, specificity and F1 scores were calculated accordingly.

3. <u>Validation against the standard image-based Caspase-3/7 assay</u>

Additionally, the AI algorithm was validated against a standard low-resolution image-based caspase-3/7 assay in a proof-of-concept toxicity study where BBB organoids were treated with staurosporine (S5921, Sigma) to induce apoptosis^27^. To that end, BBB organoids were treated with different concentrations of staurosporine for eight hours at 37°C 5% CO_2_. Half of the organoids (approx. 165) were processed to PFA-PE HE images as described above and subjected to the newly developed AI algorithm based on morphological features. The other half of the organoids (two wells per condition) were treated with Cell Event Caspase 3/7 (C10423, Invitrogen) at 2 µM for 30 minutes, according to the manufacturer’s protocol. Living organoids were then imaged at 5× (Leica Thunder 1024x1024 pixels, 14 z-stacks 27.7 µm per step, total volume 360 µm; Thunder ICC length 6.7x10^-9^, Strength 98%).

4. <u>Validation against naive organoids</u>

As a last step, the AI algorithm was validated on a separate set of 16 BBB organoids that were treated with staurosporine, as described above. Again, three blinded European board-certified veterinary anatomical pathologists evaluated and manually annotated apoptotic nuclei on a TMA HE WSI of all 16 organoids. Again, object-based accuracy was calculated as described abolve and validationn metrics included precision, recall and F1-score.

#### Quantification and statistical analysis

For whole mount immunofluorescence quantification as well as for quantification of caspase-3/7 activity in the standard image-based Caspase-3/7 assay, maximal projection images were analyzed using a custom-made automated Fiji (Fiji ImageJ 1.54) script that segments individual organoids and measures the mean fluorescence intensity of the organoid core.

## Supporting information

Supplemental Infomation

## Acknowledgements

We would like to thank Alime Teksen, Tom Albrecht, Benjamin Gutierrez Becker, Alessio Tovaglieri and Diederik van Hartog for their support in developing the AI algorithm and to our ECVP board certified pathologist colleagues Fernando Romero Palomo for annotations in BBB organoids for the AI algorithm validation. We would like to thank MicroMatrices Associates Ltd (Simon Plummer and team) for generation of the TMAs. We would like to thank Tom Leech and DLRC for their assistance in writing and formatting the manuscript.

## Author Contributions

L.B. contributed to project administration, conceptualization, data curation, methodology, investigation, visualization, writing—original draft, and writing—review & editing. C.S., E.K. contributed to conceptualization, data curation, methodology, investigation, writing—review & editing. C.Z., B.S., contributed to data curation, methodology, investigation and writing—review & editing. Ch.Z., R.N. contributed to data curation, methodology. M.T. contributed to contributed to data curation, methodology and writing—review & editing. R.V. contributed to conceptualization, supervision, writing—review & editing. A.O. contributed to writing—review & editing. N.S. contributed to project administration, supervision, conceptualization, data curation, methodology, visualization, writing—original draft, and writing—review & editing and decision to submit.

