## Supplemental Infomation for "Advanced Tissue Technologies of Blood-Brain Barrier Organoids as High Throughput Toxicity Readouts in Drug Development"

**Supplemental Information**

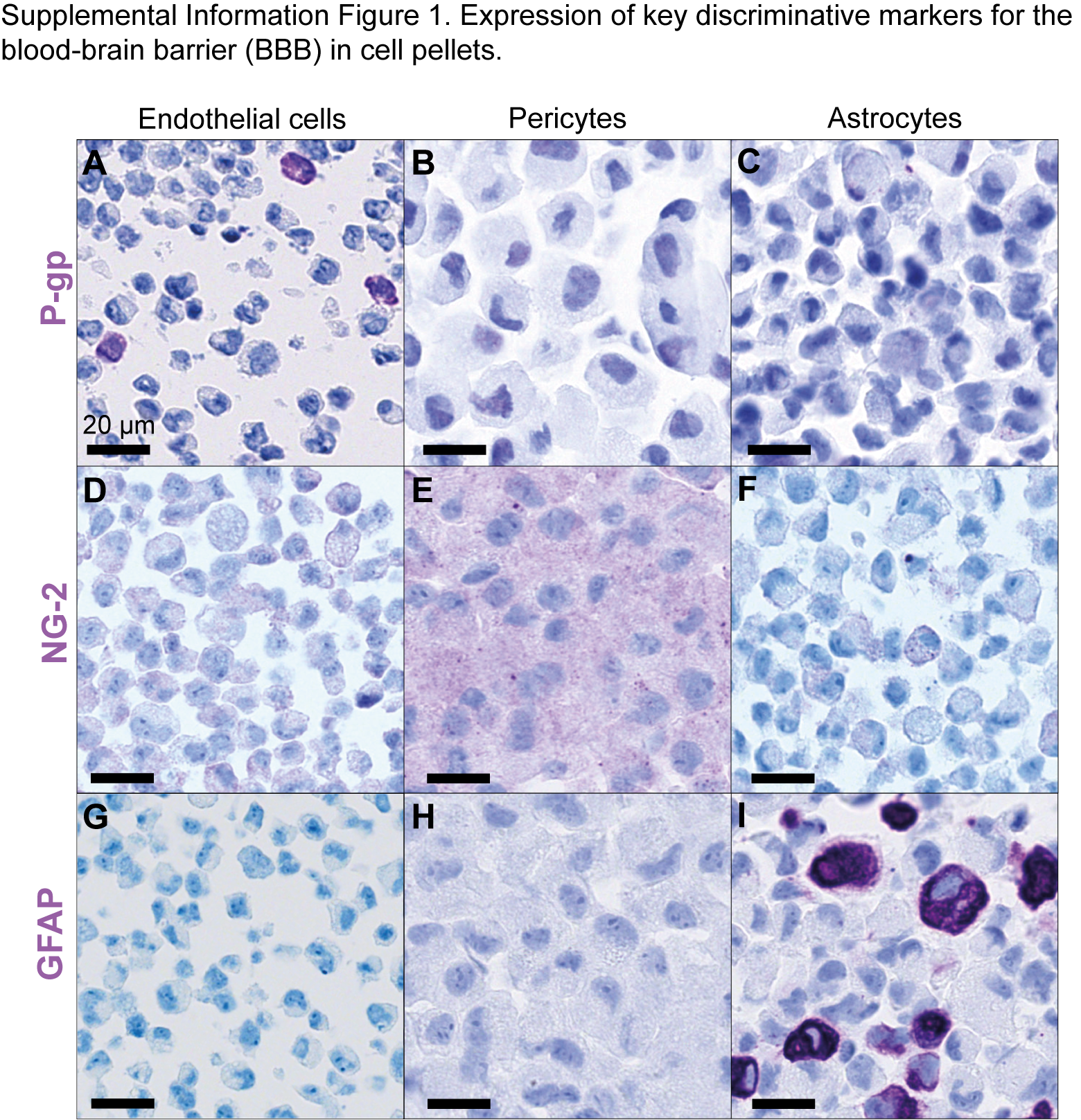

**Supplemental Information Figure 1. Expression of key discriminative markers for the blood-brain barrier (BBB) in cell pellets.** Representative image of cell pellets following immunohistochemistry. For the generation of cell pellets, adherent cells (Primary human astrocytes, ScienCell Research Laboratories; Human brain microvascular pericytes, ScienCell Research Laboratories; Human cerebral microvascular endothelial cells, hCMEC/D3, Sigma Aldrich, see Simonneau et al.^1^ for details), were collected and centrifuged in a tube. After fixation in 4% PFA for 30 minute, cell pellet was embedded in agarose and paraffin (see methods for details). IHC was performed on 4 µm thick sections on Superfrost Plus glass slides imaged at 40×. Markers were selected based on their discriminative property. Positive IHC signal in purple is variable among the different cell pellets depending on the expression pattern: P-gp positive membranous signal (purple) in endothelial cells **(A)**, while negative in pericytes and astrocytes **(B,C)**; NG-2 cytoplasmic positive signal (purple) in pericytes **(E)** while negative for endothelial cells and astrocytes **(D,F)**; GFAP cytoplasmic positive signal (purple) in astrocytes **(I)** while negative in endothelial and pericytes **(G,H)**. Nuclei are counterstained in Haematoxylin (blue). Variability of expression may be influenced by high cell passages^2^. Scale bar, 20 μm.

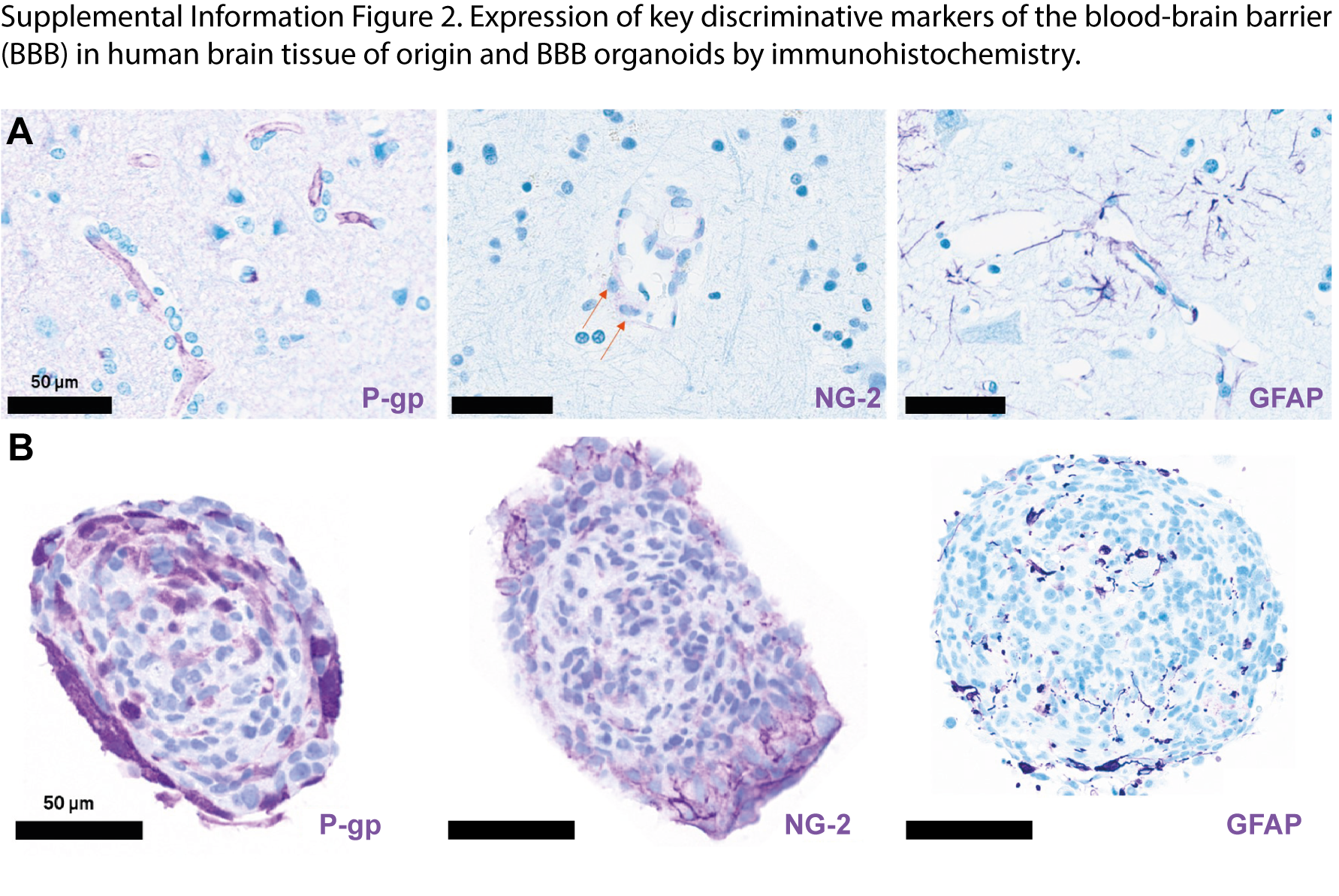

**Supplemental Information Figure 2. Expression of key discriminative markers of the blood-brain barrier (BBB) in human brain tissue of origin and BBB organoids by Immunohistochemistry.** For the generation of organoid pellets, organoids were collected and centrifuged in a tube. After fixation in 4% PFA for 30 minute, organoid pellets were embedded in agarose and paraffin (see methods for details). IHC was performed on 4 µm thick sections on Superfrost Plus glass slides imaged at 40×. **(A** and **B)** BBB-specific markers p-GP for endothelial cells, NG-2 for pericytes (orange arrows point to NG-2 positive pericytes in human cortex) and GFAP for astrocytes highlight spatial distribution of the cells contributing to the BBB in **(A)** human brain (cortex) and **(B)** BBB organoids. Nuclei are counterstained in Haematoxylin (blue). Scale bar, 50 μm.

**Supplemental Information Figure 3**. **Algorithm development to detect single apoptotic nuclei in blood-brain barrier (BBB) organoids on Haematoxylin and Eosin (HE) whole slide images.** **(A)** Objective GT generation by Caspase-3 IHC on the same slide that was previously stained for HE, using a chemical destain method. Registration of both scans allowed manual annotation of Caspase-3 positive nuclei in the HE scans (red labels) that were used for algorithm training. **(B)** Evaluation of the algorithm performance was performed by assessing the pixel-based accuracy for the objective GT by calculating the overlap of the training labels inside the algorithm output labels. **(C)** Object-based accuracy was assessed by comparing the nuclei classification (normal *vs.* apoptotic) from pathologist’s annotation (pins) to the algorithm output. Nuclei output labels are colored in bright red (positive) or dark red (negative).

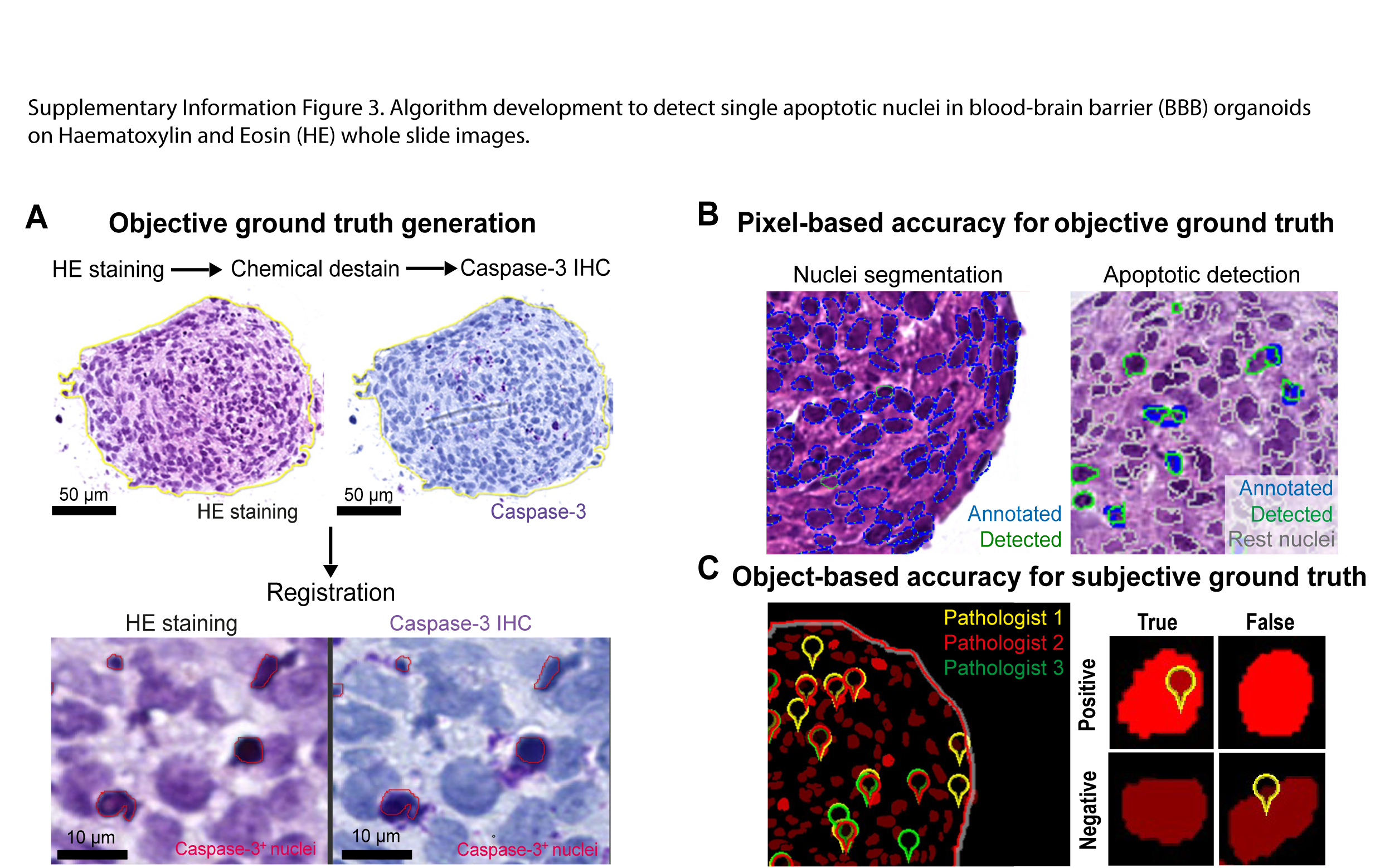

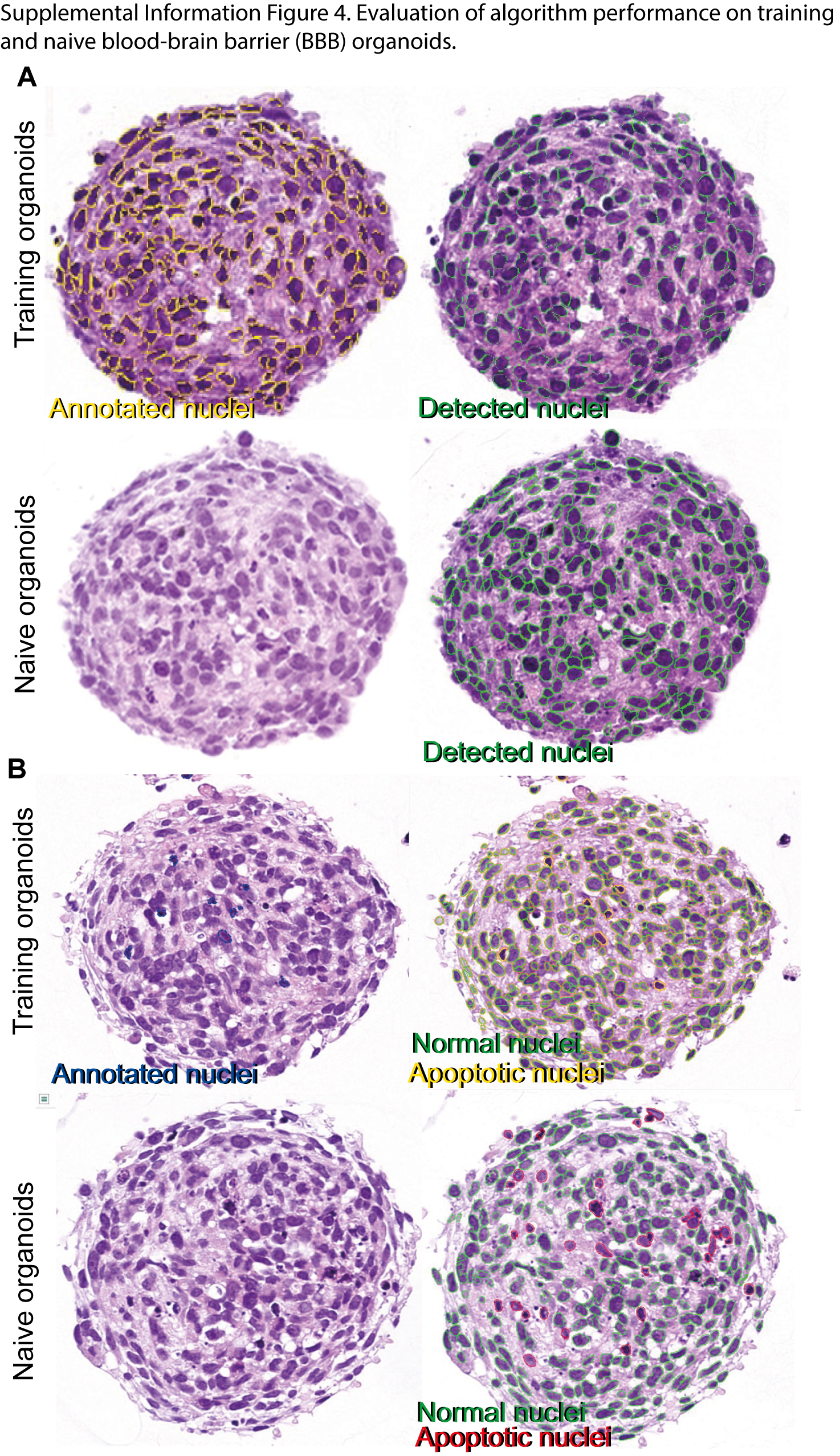

**Supplemental Information Figure 4. Evaluation of algorithm performance on training and naive organoids. (A)** Representative HE images and algorithm overlay showing the detection of nuclei by the algorithm. Manual annotations of training organoids and nuclei segmentation in training and naïve organoids. **(B)** Manual annotations of Caspase-3-positive nuclei on training organoids and algorithm overlay showing the detection of normal and apoptotic nuclei by our algorithm after training on training and naïve organoids. Pixel based accuracy was calculated by comparing the overlap of manual annotations with the algorithm output resulting in true positive, true negative, false positive and false negative pixel classification.
